# Transcriptomic and genetic analyses identify the Krüppel-like factor dar1 as a master regulator of tube-shaped long tendon development

**DOI:** 10.1101/2021.02.07.430104

**Authors:** Laurichesse Quentin, Moucaud Blandine, Jagla Krzysztof, Soler Cédric

## Abstract

To ensure locomotion and body stability, the active role of muscle contractions relies on a stereotyped muscle pattern set in place during development. This muscle patterning requires a precise assembly of the muscle fibers with the skeleton via a specialized connective tissue, the tendon. Despite evident disparities, little is known about the molecular basis of tendon diversity. Like in vertebrate limbs, *Drosophila* leg muscles make connections with specific long tendons that extend through different segments. During leg disc development, cell precursors of long tendons rearrange and collectively migrate to form a tube-shaped structure. A specific developmental program underlies this unique feature of tendon-like cells in the *Drosophila* model. We provide for the first time a transcriptomic profile of leg tendon precursors through fluorescence-based cell sorting. From promising candidates, we identified the Krüppel-like factor dar1 as a critical actor of leg tendon development. Specifically expressed in leg tendon precursors, loss of *dar1* disrupts actin-rich filopodia formation and tendon elongation. Our findings show that dar1 acts downstream of stripe as a critical regulator of cytoskeleton remodeling and mediates the recruitment of new stripe-positive tendon progenitors in a cell non-autonomous manner.

## Introduction

The musculoskeletal system comprises numerous cellular components including muscles and tendons. The assembly of these components is tightly controlled to achieve a stereotyped functional architecture. Acquisition of muscle shape and pattern thus directly relies on where the muscle fibers are anchored to the skeleton via specialized structures, the tendons. Tendons are required not only to transmit the muscle contraction force to the skeleton but also to set up a functional musculoskeletal system. The importance of coordinated development of tendons and more generally of connective tissue (CT) and muscles into an integrated system is well documented in different model organisms (Felsenthal et al., 2018; Hasson, 2011; Kardon, 2011; Kardon et al., 2003; Nassari et al., 2017; Schweitzer et al., 2010). However, the specificities of interactions between muscles and CT remain largely unknown. For example, precisely how each muscle discriminates in favor of a particular attachment site has not been clearly elucidated. Research on different models indicates that final muscle patterning relies on several elements (Hasson, 2011; Laurichesse and Soler, 2020; Schnorrer and Dickson, 2004; Valdivia et al., 2017). In *Drosophila* embryo, reprogramming muscle identity can affect the choice of attachment site (Dubois et al., 2016; Enriquez et al., 2012) and several secreted or membrane-associated proteins have been described as mediating muscle migration toward the tendon cells (Callahan et al., 1996; Kramer et al., 2001; Ordan et al., 2015; Prokop et al., 1998; Schnorrer et al., 2007; Swan et al., 2004). In avian models, transplantation studies showed that myoblast patterning was dependent on the surrounding connective tissues with Tcf-4-expressing CT cells establishing the muscle prepattern (Kardon et al., 2003, 2002; Mathew et al., 2011; Rinon et al., 2007). These observations support the hypothesis that the specificity of muscle-tendon interaction depends not only on muscle identity but also on specific CT features. In *Drosophila*, although several genes controlling muscle identity have been identified (Dobi et al., 2015), the gene code determining the identity of tendon cells remains to be characterized. Stripe/Egr-like, the key transcription factor in *Drosophila* tendon development, is a hallmark of all tendon cells (Fernandes et al., 1996; Frommer et al., 1996; Ghazi et al., 2003; Soler et al., 2004; Volk and VijayRaghavan, 1994; Vorbrüggen and Jäckle, 1997). Although a few other transcription factors are specifically expressed in tendon sub-populations such as *apterous* in wing disc-associated tendons or *GCM/Glide* in embryos, they are not considered as identity genes *per se* but are involved in subsequent tendon differentiation steps (Bernard et al., 2003; Ghazi et al., 2003, 2000; Soustelle et al., 2004). However, like vertebrates, *Drosophila* has a variety of morphologically distinct tendons. Larval monofiber muscles are linked to the exoskeleton (Frommer et al., 1996; Volk and VijayRaghavan, 1994) through a single cell attachment site, whereas large clusters of sr-positive cells specified in wing disc epithelium anchor the massive flight muscles in the adult thorax (Fernandes et al., 1996). In the fly leg, the appendicular movements are ensured by the connection between muscles and long internal tendons (Miller, 1950; Soler et al., 2004). Interestingly, these leg tendons share morphological and developmental similarities with the long tendons of the autopod in the mouse. They extend from the most distal part of the leg and elongate through the leg segments, where they are associated with extrinsic muscles (Huang, 2017; Huang et al., 2015, 2013; Soler et al., 2004; Watson et al., 2009).

Strikingly, the morphogenesis of the long tendons shares many similarities with the tubulogenesis that occurs during tracheal or salivary gland formation in *Drosophila* (Girdler and Röper, 2014; Hayashi and Kondo, 2018; Maruyama and Andrew, 2012). First, tendon cells undergo apical constriction followed by invagination without migration (Laddada et al., 2019). The cells then collectively migrate with extension of basal protrusions, leading to the elongation of a tube-shaped tendon (**Fig. 1C**). It has also been suggested that like for the developing salivary gland (Vining et al., 2005), elongating tendons could interact with surrounding tissues, especially with the myoblasts (Soler et al., 2016, 2004). We have already shown that long tendons originate from tendon precursors that are selected among the cells of leg segmental joints expressing *odd-skipped* (Laddada et al., 2019). While odd is sufficient to induce the primary invagination of the epithelial cells (Hao et al., 2003; Ibeas and Bray, 2003), stripe is required to make these cells competent to migrate and form a long tube-shaped tendon (Laddada et al., 2019). To gain a better understanding of the developmental mechanisms of appendicular tendon formation, we undertook a transcriptomic analysis of the long tendon precursors. RNA sequencing of these cells at the onset of long tendon formation enabled us to identify the associated expression of tube morphogenesis-related genes, the downregulation of which recapitulated phenotypes observed in other models of tubulogenesis. This finding functionally validates *Drosophila* leg tendon development as a new model to study tube morphogenesis.

**Fig. 1.**
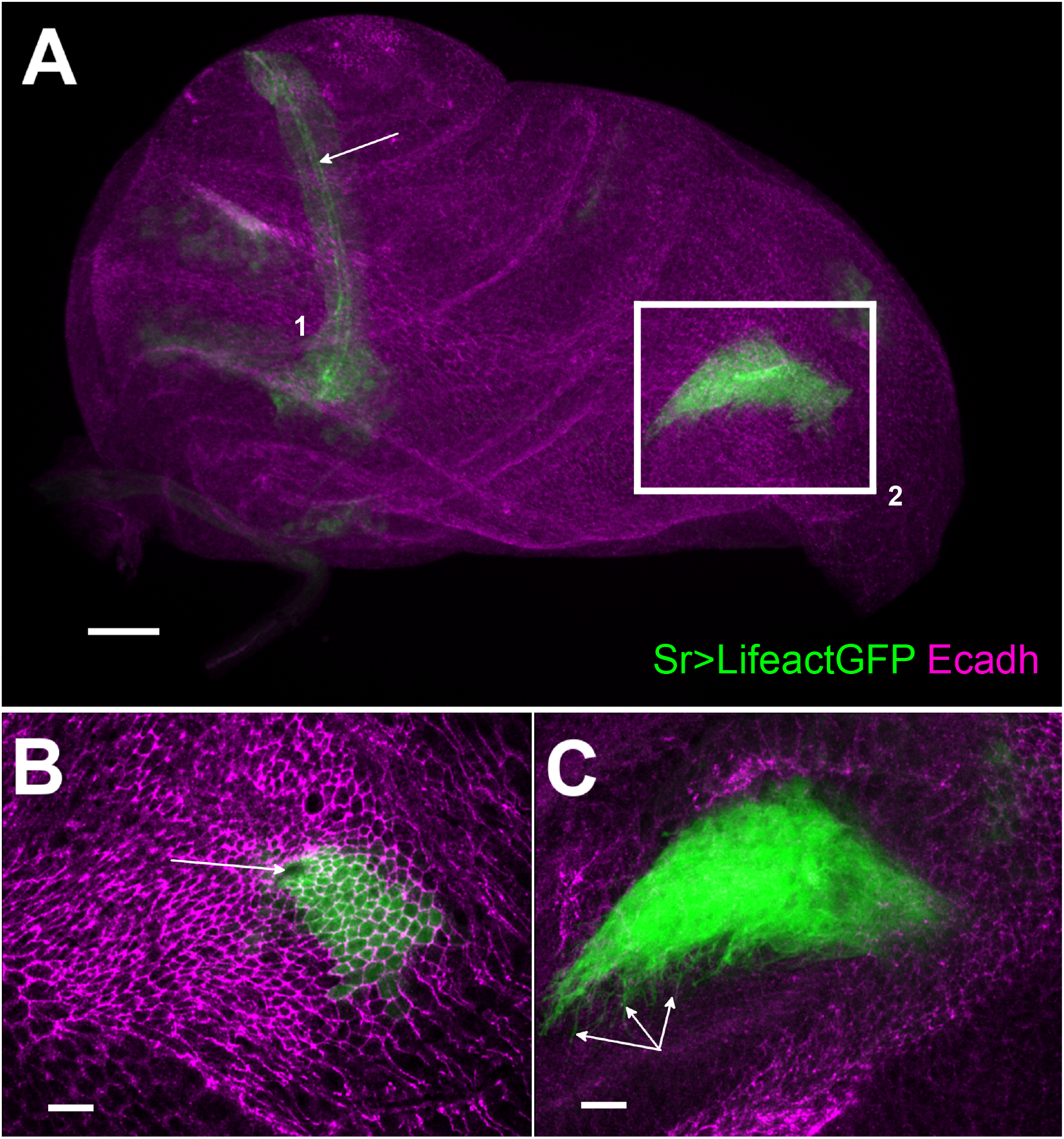
Development of long tendons from 0h APF leg disc visualized using sr-gal4 driven Lifeact.GFP expression. (A) Long tendons form from clusters of epithelial cells (stained with E-cadherin in magenta) expressing sr-gal4>UAS-Lifeact.GFP (green), long tendon (lt) of the tarsi (1) has elongated from the joint between tarsus T5 and pretarsus. Apical accumulation of Lifeact.GFP outlines the tube lumen (arrow). In the dorsal femur (box 2), the tibia levator long tendon (tilt) has started to invaginate and elongate. Scale bar 30 μm. (B) and (C) High magnification of selected confocal planes (from box 2). Scale bar 10 μm. (B) Surface view: we can observe apical constriction of Lifeact.GFP cells, revealed by E-cadherin staining, around the invaginating pit (arrow) forming the tube lumen. (C) deeper *Z* planes reveal dense arborization of actin-based filopodia at the basal side of the invaginating tendon (arrows).

We went on to perform a genetic screen to identify key transcription factors involved in the developmental program of these specific tendons. Among the positive candidates, we identified *dar1* (*dendritic arbor reduction 1*), a gene coding for a DNA-binding protein belonging to the Krüppel-like transcription factor (KLF) family. In *Drosophila*, dar1 had previously been shown to determine the multipolar morphology of post-mitotic neurons (Wang et al., 2015; Ye et al., 2011) and to downregulate the proliferation of intestinal stem cells (Wu et al., 2018). Functional analysis revealed that long tendon development was severely impaired after *dar1* knockdown. The number of cytoplasmic protrusions was drastically reduced, and the elongation, but not the invagination of tendon cell precursors was compromised. Importantly, we found that dar1 influenced tendon cell numbers without affecting the expression of *stripe*, its upstream regulator. In view of its critical effect on tendon development, we describe the role of dar1 as a key component of tube-shaped tendon morphogenesis, which could act in turn on *stripe* cell specification in a non-autonomous manner.

## Results

### Transcriptomic profiling of appendicular long tendon precursors reveals a gene expression signature of tubulogenesis

We had previously shown that precursors of the long tendons are clusters of epithelial cells selected among the cells of leg segmental joints. The Notch pathway initiates *stripe* expression, the key factor in tendon cell differentiation, in the leg tendon precursors (Laddada et al., 2019; Soler et al., 2004).

These sr-positive cells invaginate and migrate to form the tube-shaped long tendon between the end of the larval stages and the first hours of pupal formation. To gain a better understanding of the molecular mechanisms underlying these dramatic cell morphological changes, we undertook a whole transcriptome RNA-seq analysis of these cells at the onset of pupation (0h APF). Total RNAs were prepared from FACS-isolated sr-gal4>UAS-GFP cells from 0h APF leg discs and processed for whole-transcriptome RNA-seq, followed by bioinformatic analysis (for cell-sorting and RNAseq detailed protocols and validations see Materials and Methods and Supplementary Material). With the FPKM cut-off at 10 for reliable detection of gene expression, we found that a wide array of genes (5479) were significantly expressed in sr-gal4>UAS-GFP cells, probably reflecting the developmental plasticity exhibited by *Drosophila* imaginal disc cells (McClure and Schubiger, 2007). However, gene ontology (GO) analysis of these genes showed an over-representation of the GO terms related to “locomotion” and “muscle structure development” with numerous genes known to be involved in muscle attachment site development and/or the formation of myotendinous junctions (**Table S1**). Considering GOs related to pathways, we found an over-representation of Notch, Wnt and MAPK pathways, consistent with our previous results and those of others, showing that these pathways play a key role in tendon development in *Drosophila* (Laddada et al., 2019; Lahaye et al., 2012; Soler et al., 2004; Yarnitzky et al., 1997). Finally, this analysis also pointed to GO terms relative to tube morphogenesis such as “epithelial tube morphogenesis” or “regulation of tube size”, supporting our earlier claim that long tendon development shared common features with salivary gland or tracheal tube morphogenesis (Girdler and Röper, 2014; Hayashi and Kondo, 2018; Laddada et al., 2019). To functionally validate this point, we crossed the sr-gal4 line with UAS-RNAi lines directed against genes known to regulate tube morphogenesis. First, we tested genes known for their role in many aspects of cell adhesion, migration or maintenance of tissue integrity during morphogenetic processes such as tubulogenesis. As expected, the expression of RNAi against genes such as the βPS integrin totally disrupted long tendon formation (data not shown). More interestingly, knocking down the expression of genes controlling the length of developing tubes (Kerman et al., 2008; Luschnig et al., 2006) such as *serpentine, vermiform* or *lolal*, led to an excessive elongation of the long tendons in the mature leg (**Fig. S1**). All in all, these results validate our approach to identifying new regulators of appendicular long tendons and support a pivotal role of tubulogenesis in their development.

### Deciphering the core genetic program of leg tendon development

Because transcription factors are fundamental in controlling the developmental program to build any structure during development, we carried out a lethality and climbing-based *in vivo* RNAi screen targeting genes encoding for TFs that we found specifically enriched in our RNAseq data. Remarkably, of the 5479 genes with FPKM>10, transcripts of 31 genes encoding for predicted transcription factors (Rhee et al., 2014) exhibited more than 1.5-fold higher levels of expression in the GFP+ cells than in the whole leg disc cells (input), strongly suggesting that these TFs have a relevant specific role in the development of the leg long tendon (**Table S2**). We performed this RNAi screen by crossing tendon-specific UAS-Dicer2;sr-gal4 line with UAS-RNAi lines targeting these 31 candidates from two independent libraries: TRiP (Bloomington Stock Center) and VDRC (Vienna Stock Center). To limit the number of false positives due to off-target effects, we did not include UAS-line RNAi from the VDRC, for which more than two off-targets were predicted. In this way, we crossed 53 UAS-RNAi lines to target the 31 selected genes, and we could test two different lines for 71% of them (22/31) (**Table S2**). F1 generation were screened for developmental lethality and/or climbing defect (for the detailed screen protocol, see Materials and Methods). Of the 31 candidates analyzed, 7 showed a significant embryonic (or early larval stage) lethality, with more than 20% of embryos unable to hatch or dying before their third larval stage, 12 displayed a pupal lethality score higher than 20%, and 14 showed a climbing defect. In all, nearly 65% (20/31, including *stripe* itself) of the candidate genes were positive for at least one readout and for eight of them this result was obtained with two different RNAi lines (**Table 1**). Of these positive hits, we found *odd* and *drm* both members of the *odd-skipped* gene family that act downstream of the Notch pathway to promote leg tendon growth (Laddada et al., 2019). We also identified *esg*, which interferes with the Notch pathway during stem cell maintenance/differentiation (Korzelius et al., 2014), and which is required during tracheal morphogenesis (Miao and Hayashi, 2016). These positive genes strengthen our strategy to identify new regulatory factors of leg long tendon development. Other candidates, including six uncharacterized predicted transcription factors (CG numbers), have never been reported or investigated in tendon development.

**Table 1.** List of positive candidate genes based on RNAi screen.

| Gene | Known/predicted function | Lethality | Locomotion Defect | origin of RNAi line showing phenotype(s) | FC value | Human Ortholog# |
| --- | --- | --- | --- | --- | --- | --- |
| **Drm** | imaginal disc-derived leg joint morphogenesis, tube morphogenesis | metamorphosis | YES | VDRC | 4,1 | OSR1/2 |
| **Dar1** | positive regulation of dendrite morphogenesis | metamorphosis | YES | TRiP and VDRC | 4,7 | KLF5/KLF4 |
| **CG7785** | - | - | YES | TRiP and VDRC | 5 | SPRYD7 |
| **CG11317** | - | embryo or early larval stages (100%) | nd | TRiP* | 1,8 | - |
| **Ato** | eye-antennal disc morphogenesis, sensory organ precursor cell fate determination | - | YES | TRiP | 5,5 | ATOH7 |
| **Esg** | stem cell homeostasis, tracheal morphogenesis, Notch pathway... | metamorphosis | YES | TRiP* | 1,8 | SNAI1/2 |
| **Odd** | imaginal disc-derived leg joint morphogenesis, tube morphogenesis | metamorphosis | YES | VDRC | 2,42 | OSR1/2 |
| **NF-YB** | R7 cell differentiation, regulation of MAPK cascade | - | YES | TRiP and VDRC | 1,6 | NFYB |
| **e(y)2** | nuclear export of mRNA, SAGA complex | metamorphosis/embryo or early larval stages | no | TRiP | 1,7 | ENY2 |
| **CG15435** | unknown function | metamorphosis/embryo or early larval stages | YES | TRiP | 1,5 | - |
| **REPTOR-BP** | TORC1 signaling, response to starvation | metamorphosis | YES | TRiP | 1,7 | CREBL2 |
| **CG6276** | unknown function | metamorphosis | YES | TRiP and VDRC | 1,6 | - |
| **Sox15** | wing disc development, apoptotic process | - | YES | TRiP and VDRC | 1,6 | SOX18 |
| **CG9948** | unknown function | - | YES | VDRC* | 1,5 | - |
| **CG9650** | unknown function | metamorphosis | YES | TRiP and VDRC | 1,6 | BCL11A/BCL11B |
| **Dichaete** | Axon guidance, hindgut development | embryo or early larval stages | YES | VDRC | 1,6 | SOX21 |
| **Dlip1** | unknown function | embryo or early larval stages | no | TRiP | 1,91 | - |
| **E(spl)my-HLH** | Notch pathway, neuroblasts proliferation | metamorphosis | no | TRiP | 2,11 | HES2 |
| **l(2)k10201** | unknown function | metamorphosis/embryo or early larval stages | no | TRiP and VDRC | 2,09 | ZFP511 |
| **stripe** | tendon cell fate induction | embryo or early larval stages (100%) | nd | TRiP and VDRC | 7,89 | EGR |

\* no alternative (VDRC or TRiP) available stock

We then considered the vertebrate orthologs of these newly-highlighted TFs to identify functional evolutionary conservation. We note that of the 14 conserved genes, the orthologs of *dar1* (KLF5 and KLF4) and *CG9650* (BCL11A) were shown to be differentially regulated in limb tendon cells during mouse development in two independent studies (Havis et al., 2014; Liu et al., 2015). Strikingly, for each of these genes, the expression of two different RNAi lines gave significant adult climbing defect and/or metamorphosis but no embryonic lethality (**Table S2**). These observations strongly suggest that these two genes are likely candidates for the specific development of appendicular long tendons. Below we analyze the expression and function of *dar1* characterized by high transcript enrichment in our RNAseq datasets.

### *dar1* is expressed in leg tendons but not in embryonic and flight muscle attachment sites

To confirm the expression of *dar1* in the leg disc tendon, we performed immunostaining on sr-gal4>UASmcherryNLS leg discs from third larval instar to 5h after pupal formation (APF) using dar1 antibody (Ye et al., 2011). *dar1* expression colocalizes with *stripe* expression domains, within the cells corresponding to the specified long tendons of the leg (**Fig. 2**). *dar1* was first found in the two earliest clusters of sr-positive cells that would form the long tendon of the tarsi (lt) and in tibia levator tendon (tilt) in the dorsal femur (**Fig. 2 A,B**). Subsequently, during early pupation this expression was maintained in lt and tilt and new sr-positive clusters corresponding to other long tendons in different leg segments started to be specified (**Fig. 2C-F**). Importantly, *dar1* expression was observed in all these clusters and correlated with our RNA-sequencing data generated from tendon precursors at early pupal stage (0–2h APF), which showed a high specific enrichment of dar1 expression in sr-gal4>GFP cells compared to IP (FC=4.7), strengthening the reliability of our data. To determine whether *dar1* was a hallmark of all tendon cells in *Drosophila*, we also analyzed its expression in embryos and in wing discs (**Fig. S2**). In embryos, we could not detect dar1 in sr-gal4>UASmcherryNLS positive tendon cells (**Fig. S2A**,**B**). Likewise, tendon precursors of flight muscles in the wing disc did not show any *dar1* expression (**Fig. S2C**,**D**).

**Fig. 2.**
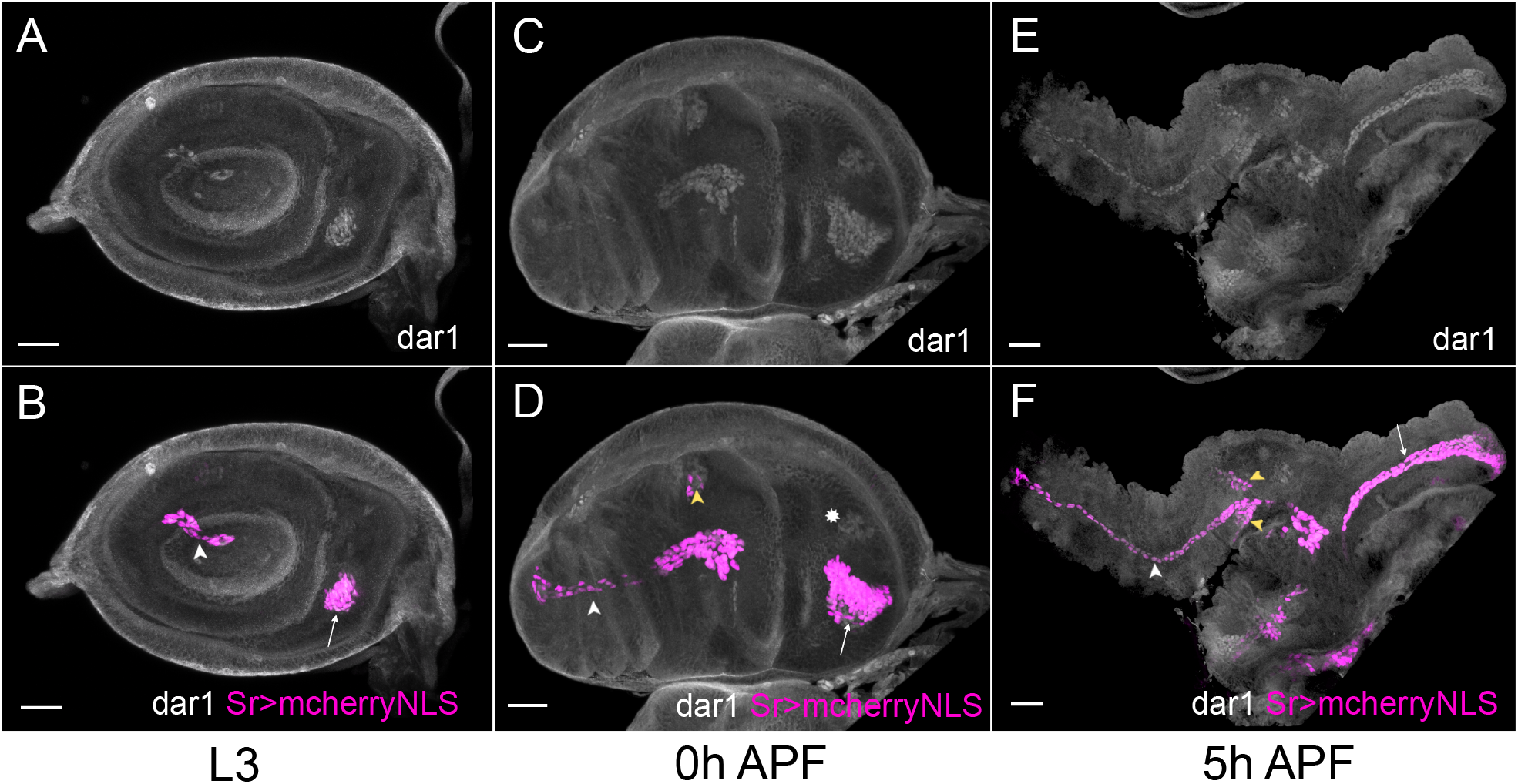
*dar1* expression pattern co-localizes with *stripe* expression in *Drosophila* leg disc. Selected optical sections of sr-gal4>UAS-mcherryNLS (magenta) leg discs immunostained with anti-dar1 (gray) at different steps of development. (A-B) In L3 leg disc, *dar1* and *mcherryNLS* expressions colocalize in cells prefiguring the future lt in the tarsi (white arrowhead) and the future tilt in the dorsal femur (arrow). This co-expression is maintained at 0h APF (C-D) and 5h APF (E-F) in elongating tendons, whereas newly-specified sr>mcherry-positive cells, in tibia segment notably, also start to express *dar1* (yellow arrowheads). Note that *dar1* is also expressed at a low level in the apparent chordotonal organ at 0h APF (asterisk). Scale bar 30µm.

All in all, these results strongly suggest that dar1 plays a specific and critical role in the establishment of these unique long tendons in *Drosophila*.

### Correct adult muscle patterning required *dar1* tendon expression

To determine whether the adult locomotion defects observed in tendon-specific *dar1* KD could be explained by an abnormal development of the appendicular musculotendinous system, we analyzed adult leg myotendinous architecture of sr-gal4>UAS-Lifeact.GFP, UAS-dar1RNAi flies combined with heterozygous null *dar1*^*3010*^ allele to potentialize dar1RNAi efficiency. As shown by longitudinal cryosections along the proximo-distal axis of the sr-gal4,dar1^3010/+^>UAS-Lifeact.GFP, UAS dar1RNAi legs, long tendons in all segments were often shortened or even missing, and so muscle pattern was strongly affected (**Fig. 3**). Major muscles, labeled by phalloidin, in control legs displayed a feather-like pattern with each fiber attached on one side to a long tendon and on the other to an individual cuticle muscle attachment site (cMAS). By contrast, when long internal tendons were affected, both ends of a muscle fiber were anchored to a cMAS, generating a transverse fiber. This is clearly visible in the ventral tibia where the long tendon appears much shorter in the visualized cryosections (**Fig. 3E**). Thus in the proximal region of this segment, both ends of the muscle fibers are attached to cMAS, whereas in the distal part, the fibers still connect to the shortened remaining ventral long tendon (Fig. 3F). We could also observe a highly disorganized muscle pattern in the dorsal femur that coincided with no visible tilt in this cryosection. Our observations thus indicate that when *dar1* expression is specifically lowered in leg tendon precursors, resulting long tendons are severely affected, leading to abnormal muscle patterning. Moreover, even in the absence of long tendons, the muscle fibers can still attach to cuticle via cMAS, suggesting that the latter, like those of flight and embryonic muscles, do not required dar1 activity.

**Fig. 3.**
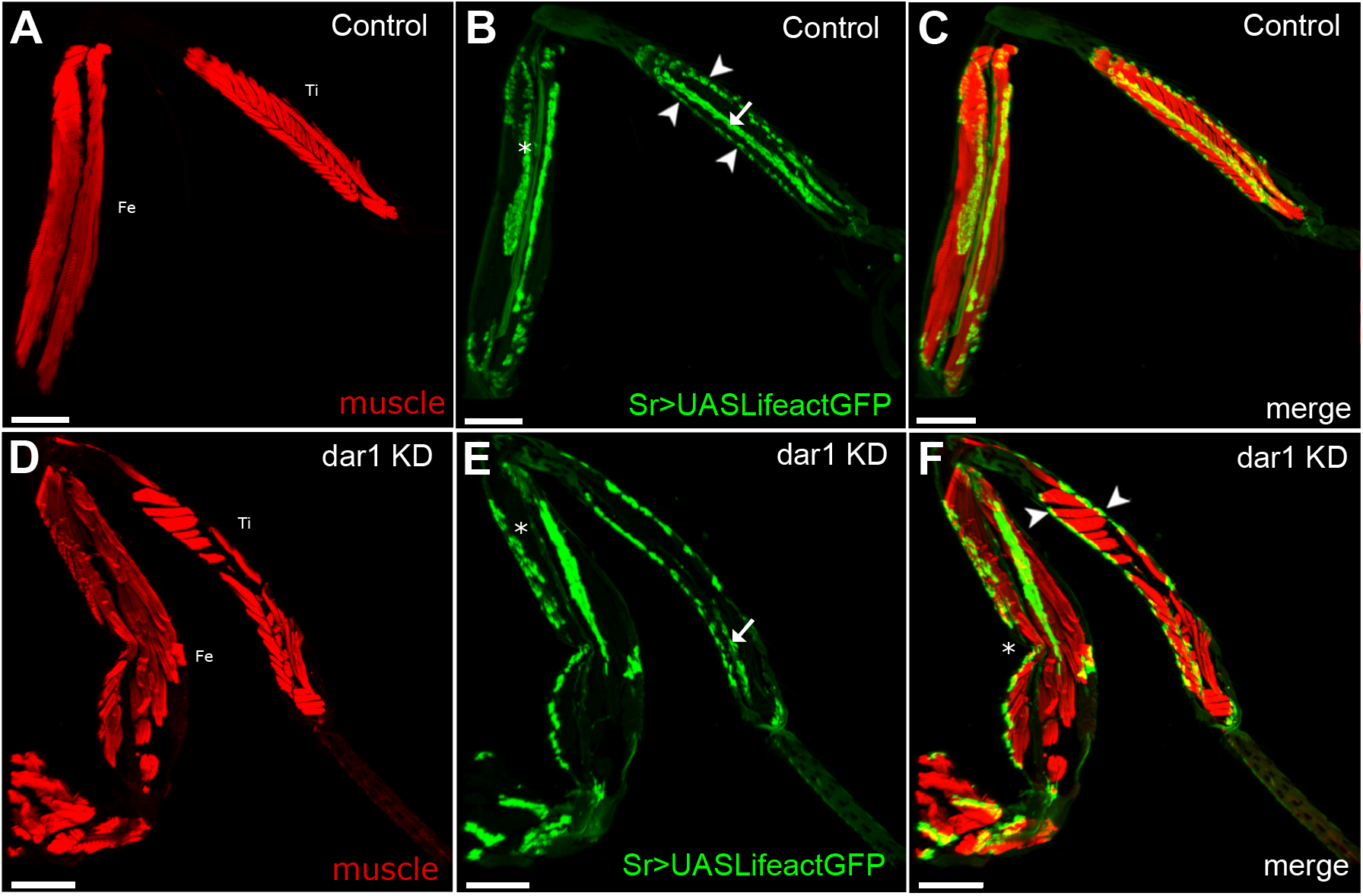
Knockdown of *dar1* expression in long tendons alters myotendinous architecture of adult leg. Combination of confocal images from adult leg cryosections (Tibia: Ti, Femur: Fe); muscles are stained with stained phalloidin (red). Tendons are visualized with Lifeact.GFP (green). (AC) In each segment of control sr-gal4,dar1^3010/+^>UAS-Lifeact.GFP legs, major muscles display a feather-like pattern. For instance, fibers of the muscle in the ventral tibia are attached on one side to the tarsi depressor long tendon (arrow in B) and on the other to an individual cuticle muscle attachment site (arrowheads in B). (D-F) on this cryosection of sr-gal4, dar1^3010/+^>UASLifeact. GFP,UAS-dar1RNAi leg, the tilt in the dorsal femur is absent (asterisk in E compared to B) and the long ventral tendon appears shorter in the tibia (arrow in B and E). Alteration or absence of these long tendons leads to the misattachments of muscle fibers with both ends attached to cuticular attachment sites (arrowhead in F) leading to the formation of transversal fibers. Note that disruption of myotendinous architecture seems to affect the morphology of the leg, with a pinch in the femur segment (asterisk in F). Scale bar 100μm.

### dar1 is required for tubulogenesis-like morphogenesis of leg tendons

To better characterize the role of dar1 in long tendon development, we focused our analysis on the time when tendon precursors started to express *dar1*, from the L3 larval stage to the early hours of pupation. We specifically analyzed the tarsal lt, extending from pre-tarsus to femur, and the tilt in the dorsal femur. These two tendons form earliest: they are specified at the early third larval instar and then invaginate and elongate until the first hours of metamorphosis (Laddada et al., 2019; Soler et al., 2004). Tendon specific depletion of *dar1* led to apparent shortening of these elongated structures in early metamorphosis (**Fig. 4A-F**). To quantify this phenotype, we measured the relative size of the lt and tilt (for measurement details see Materials and Methods) at three developmental time points (late L3, 2h and 4h APF) in control (sr-gal4,dar1^3010/+>^UASLifeact.GFP) and *dar1* KD (sr-gal4,dar1^3010^/+>UAS-Lifeact.GFP, UASdar1RNAi) leg discs. For each of these developing tendons, we observed a significant reduction of their length at the three different time points when *dar1* expression was lowered (**Fig. 4G-H**). The tarsal tendon was very significantly shortened as early as late L3, whereas the difference in size of the tilt in dorsal femur was more pronounced at later stages (4h APF). This is probably because tarsal lt is specified and starts to elongate slightly earlier than the one in the dorsal femur (Soler et al., 2004) and is therefore longer at a given time point. Thus, the length difference compared with control was greater in earlier stages for the tarsal lt than for the tilt in dorsal femur. We infer that the apparent smaller size of the tendons after *dar1* KD becomes more and more significant as the tendons elongate, supporting a potential function of dar1 in this process. Likewise, 2h APF leg discs from rare escapers carrying homozygote *dar1*^*3010*^ null mutation, stained against discs large (dlg) septate junction marker, showed no visible elongating structure (**Fig. 5A,B**). Remarkably, in the location of the most distal tarsal segment where lt normally starts to develop, we could still see an accumulation of dlg protein prefiguring the local folding of the epithelium and the lumen formation (**Fig. 5A,B**). These observations indicate that dar1 is not needed for initial steps of epithelium invagination but is then required for the tube-like tendon elongation.

**Fig. 4.**
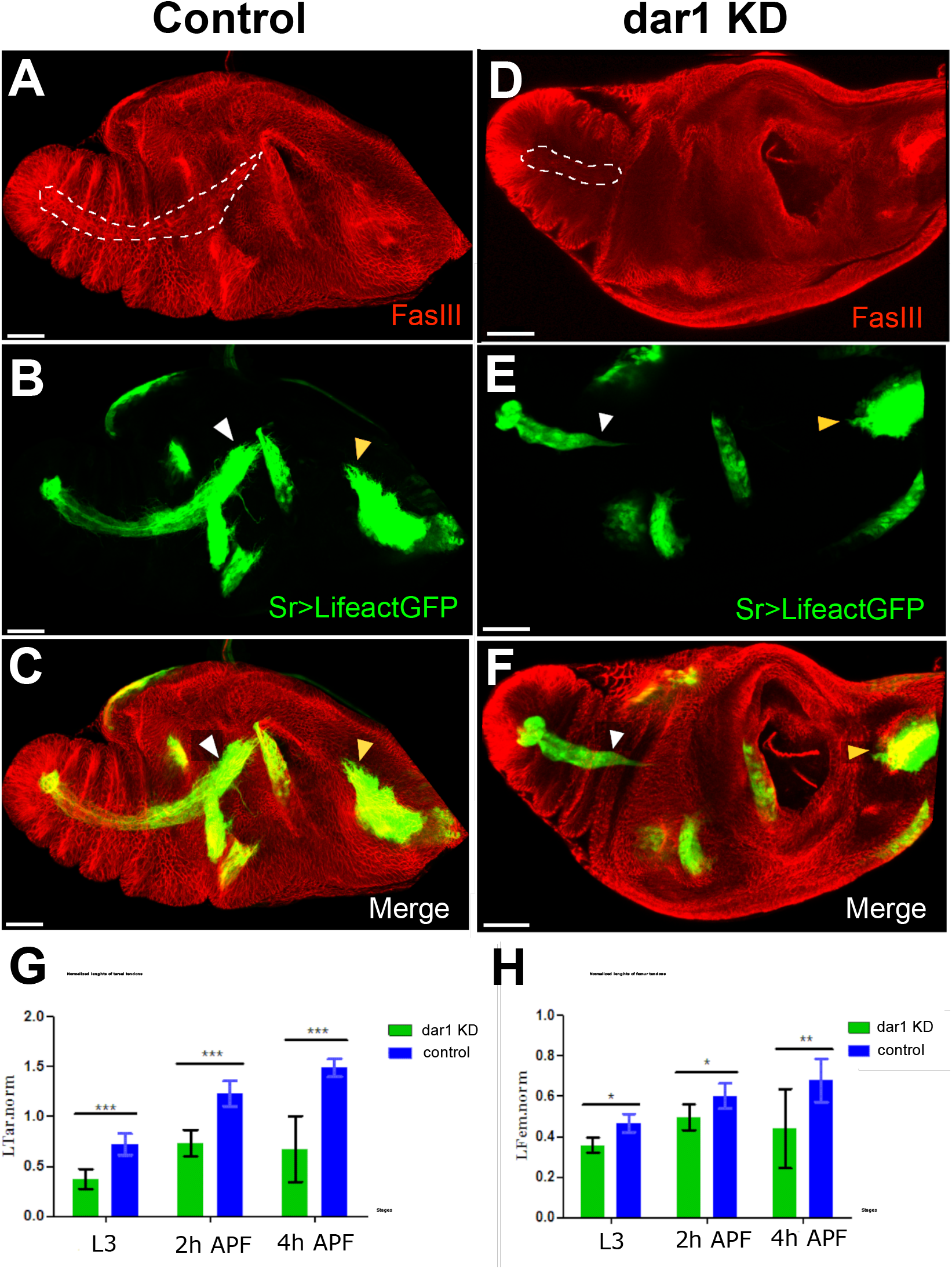
dar1 is required for the elongation of long tendons. Confocal images of sr-gal4,dar1^3010/+^>UAS-Lifeact.GFP (A-C) and sr-gal4,dar1^3010/+^>UAS-Lifeact.GFP,UAS-dar1RNAi (D-F) leg discs at 3h APF immunostained with anti-fasIII. On this optical section, the long elongating tendon of the tarsi is revealed by UAS-Lifeact.GFP and FasIII (dashed line A and D). In *dar1* KD leg disc (D-F), the tarsal lt appears much shorter than in the control (white arrowheads in B,C and E,F). At this stage of development, other tendons are less elongated than the one in the tarsi and therefore their difference in length between control and *dar1* KD is not as obvious, except for the tilt in the dorsal femur (yellow arrowhead in B,C and E,F), which also appears shorter. (G-H) Graphs showing the lengths of the lt in tarsi (G) and tilt femur (H) in control leg discs versus and *dar1* KD leg disc at three time points: late L3 (control *n*=11, *dar1* KD *n*=11), 2h APF (control *n*=10, *dar1* KD *n=*17) and 4hAPF (control *n=*5, *dar1* KD *n=*5). Scale bar 40µm.

**Fig. 5.**
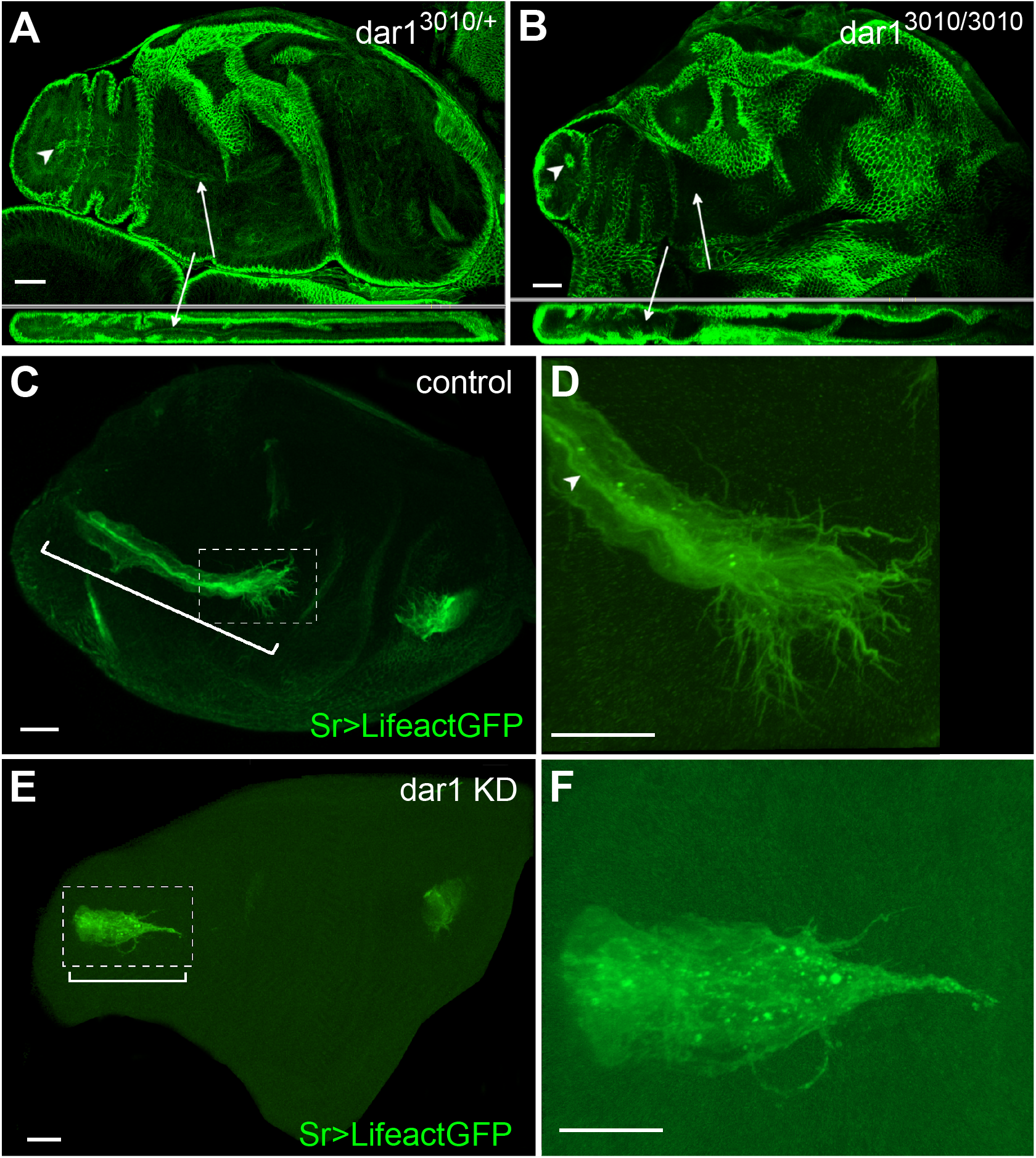
dar1 is not required for the epithelium invagination but is needed for actin-rich filopodia arborization of long tendons. (A-B) Confocal sections of 2h APF leg discs immunostained with anti-dlg. (A) selected optical sections of control *dar1*^*3010/+*^ heterozygous leg disc. dlg accumulation reveals lumen aperture on the most distal tarsal segment (arrowhead) and also stained a long internal structure corresponding to tarsal lt (arrow). (B) In leg disc of rare *dar1*^*3010/3010*^ homozygous mutant escapers, local constriction of the epithelium still underlines the local invagination of the epithelial cells, but no elongating internal structure is formed (arrows). (C-F) Confocal sections of 0h APF leg discs from sr-gal4,dar1^3010/+^>UAS-Lifeact.GFP and sr-gal4,dar1^3010/+^>UAS-Lifeact.GFP,UAS-dar1RNAi 2h APF pupae. On these selected optical sections, in *dar1* KD condition tarsal lt (bracket in E) appears much shorter than in the control (bracket in C). (D) and (F) are higher magnifications from (C) and (E) respectively. (D) In the control leg disc, leading cells at the distal part of the elongating tendon display numerous actin-rich protrusions at their basal membrane and apical accumulation of actin underlines the tube lumen (arrowhead). (F) In the *dar1* KD leg disc, filopodia arborization is systematically affected with a marked decrease in protrusion number (number of samples, control *n*=10, *dar1* KD *n*=17). Overall Lifeact.GFP distribution is strongly affected and tube lumen cannot be distinguished in *dar1* KD. Scale bar 20 µm

Besides this elongation defect, *dar1* RNAi expression in long tendon precursors results in a severe depletion of filopodia as revealed by UAS-Lifeact.GFP expression (**Fig.5 C-F)**. For instance, the total number of filopodia at the tip of the tarsal lt in control 2h APF disc was systematically greater than 30 (*n*=10), whereas in *dar1* KD leg discs, tarsal lt systematically displayed fewer than five remaining protrusions (*n=*17). This result indicates that the reduction of *dar1* expression directly or indirectly affects the formation of actin-rich filopodia. Moreover, while the tendon lumen in control leg discs is underlined by apical Lifeact.GFP accumulation, in *dar1* KD leg disc we observed a fluorescent dotty pattern reflecting a general actin polymerization defect.

Taken together, our results show that dar1 participates in the cellular events required for actin cytoskeleton organization and specifically filopodia formation. Filopodia are dynamic actin-based projections with sensing capacity, thought to promote cell motility (Jacquemet et al., 2015). *dar1* downregulation thus leads to a tendon elongation defect, which could be attributed to a defect in the ability of tendon cells to migrate.

### dar1 regulates the number of Stripe cells non-autonomously

➢ *dar1* depletion induces loss of sr-positive progenitors without affecting proliferation nor inducing apoptosis

Although it has been shown that the number of cells that make up a tube is not closely correlated to tube length (Beitel and Krasnow, 2000), the question arose of whether the total number of sr-gal4 positive cell was affected after *dar1* KD. To find out, we counted the number of cells constituting both tarsal and dorsal femur tendons in sr-gal4,dar^3010/+^>UASmcherryNLS control pupae, and in sr-gal4,dar^3010/+^>UASmcherryNLS, UASdar1RNAi pupae at early stages of metamorphosis (**Fig. 6A-C**). First, in control leg disc we saw an increasing number of sr-gal4>mcherryNLS cells from 0h to 5h APF for both tendons, indicating that during the elongation process new sr-positive cells were recruited. Next, we found that the number of sr-gal4>mcherryNLS cells was significantly reduced in *dar1* KD leg discs at the two different time points compared to the control leg disc. (**Fig. 6A-C)**. Lastly, in *dar1* depleted tarsal tendon, the number of sr-gal4>mcherryNLS cells was not statistically different between 0hAPF and 5hAPF, suggesting that no further tendon cells were recruited between these two time points, unlike in controls. Our results thus show that elongation of long tendon defects observed on dar1 downregulation correlates with a default of sr-gal4>mcherryNLS cell numbers. As a decrease in the number of cells may be due to a default in cell proliferation and/or to an increase in cell death, we tested these two variables in sr-gal4,dar^3010/+^>UASmcherryNLS, UASdar1RNAi leg discs. As tendon cells are most likely post-mitotic cells and because dar1 has been shown to restrict the proliferation of intestinal stem cells (Wu et al., 2018), we did not favor a default of proliferation, but we immunostained *dar1* KD and control leg discs using phospho-histone 3 antibody. Although numerous leg disc cells proliferated from mid-L3 stage (beginning of tendon cell specification) to early metamorphosis, only rare mitotic events could be observed among sr-gal4>mcherryNLS cells in this time window in both *dar1* KD and control conditions (**Fig. S3**).

**Fig. 6.**
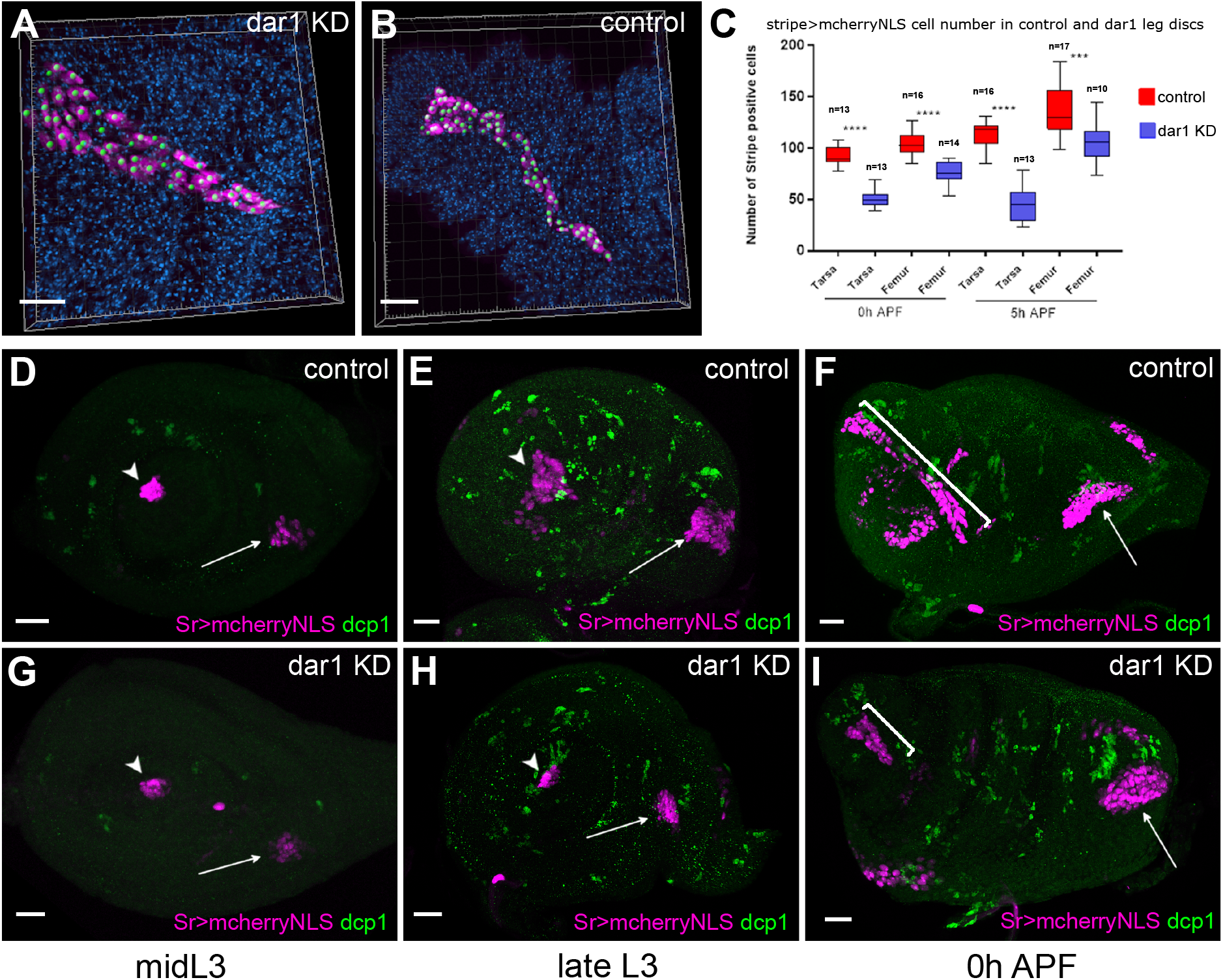
Reduction of tendon cell number in *dar1* KD leg discs is not due to ectopic apoptosis. (A-B) examples of semi-automated counting tendon cells (magenta) from Z-stack projections of confocal images of lt from sr-gal4,dar1^3010/+^>UAS-mcherryNLS, UAS-dar1RNAi (A) and sr-gal4,dar1^3010/+^>UAS-mcherryNLS (B) 5h APF leg discs, stained with DAPI (blue). Each green spot corresponds to a single cell. (C) Box-plot diagram comparing number of mcherry-positive cells in lt and in the tilt, in control and in *dar1* KD leg discs at two different time points. Box boundaries and horizontal bar indicate 25/75 percentile and mean value, respectively. (D-I) Confocal images of sr-gal4,dar1^3010/+^>UAS-mcherryNLS (magenta) (D-F) and sr-gal4,dar1^3010/+^>UAS-mcherryNLS,UAS-dar1RNAi (G-I) leg discs immunostained with anti-dcp1 (green) at different times of development. Arrowheads and arrows point to tarsal lt and tilt in femur, respectively. In both control (D) and *dar1* KD (G) L3 leg discs, cell death cannot be detected in lt or tilt. At late L3, several apoptotic cells are observed throughout the leg discs (E, H) but none of them are tendon cells. As stated previously, at this time there are already fewer tendon cells in the *dar1* KD leg disc (H) than in the control (E). At early metamorphosis, lt (bracket) appears much shorter in the *dar1* KD leg disc (I) than in the control (F), but no specific cell death is observed in this tendon. Observed discs: control *n*=19, *dar1* KD, *n*=21. Scale bar 20µm.

To determine whether loss of sr-gal4>UASmcherryNLS cells could result from cell death increase, we performed immunostaining against caspase-activated dcp1 (**Fig. 6D-I**). Between the time of specification of first sr-positive cells and the early step of metamorphosis, we could not detect any sr-gal4>mcherryNLS apoptotic cells in *dar1* KD leg discs, while the default in cell number was evident as early as the late L3 stage (compare **Fig. 6 E,H**). We confirmed this result by expressing GC3Ai fluorescent apoptotic sensor, which efficiently follows apoptotic cell dynamics tissue-specifically (Schott et al., 2017). Thus, when GC3Ai apoptotic sensor was specifically expressed in sr-gal4 cells, rare events of apoptosis were occasionally observed in both control and *dar1* KD leg discs (**Fig. S4)**. We concluded that the up to 50% fewer tendon cells in *dar1* KD context could not be explained by any marked increase in cell death or a lower proliferation rate.

➢ Epistatic relationship between *dar1* and *stripe* expressions

One obvious cause of missing tendon cells in the *dar1* KD leg disc would be a positive requirement of dar1 to induce and/or maintain *stripe* expression (and so sr-gal4 expression). Because all the phenotypes we observed were obtained using sr-gal4 driver to induce UAS-dar1RNAi, a cell-autonomous requirement of dar1 in inducing *stripe* expression is very unlikely. However, we tested this possibility by monitoring *stripe* expression in *dar1*^*3010*^ homozygous null-mutant escapers at the time when initial pools of *stripe* expressing cells were set up in tarsi and dorsal femur (between early and mid-L3 larval stage). As expected, stripe was still clearly detected at the L3 stage in both clusters in dar1^3010h^eterozygous or homozygous mutants, indicating that *stripe* initial induction was dar1-independent **(Fig. 7A-D)**. Conversely, expression of *dar1* was completely abolished when a dominant negative form of stripe was expressed in the leg disc epithelium (**Fig. S5)**. These results strongly suggest that stripe acts upstream to regulate *dar1* expression in the developing tendon, and that dar1 is then needed to correctly pattern long tendons with the right number of sr-positive cells. Finally, to determine whether the lack of sr-positive cells in sr-gal4,dar^3010/+^>UASdar1RNAi context could be explained by a role for dar1 in long-term maintenance of *stripe* expression, we performed a G-TRACE experimental analysis (Evans et al., 2009). This technique is used to determine the expression of a gal4 driver (here sr-gal4) at a current time point by inducing UAS-RFP expression **(Fig. 7F,I**), and its historical expression by inducing UAS-Flipase expression. Flp enzyme recognizes FRT sites and removes the STOP cassette in the Act-FRT-STOP-FRT-GFP construct, allowing a permanent expression of GFP. Accordingly, if any cell has been committed as a sr-gal4 positive cell at any time during development and then loses this expression, it should still be detectable through sustainable GFP expression. In other words, RFP expression reveals real-time gal4 expression and GFP expression its past expression. In this way, we could see in both *dar1* KD and control 5h APF leg disc, cells that currently expressed sr-gal4 (RFP+) are also GFP+ **(Fig. 7E**,**H)**. This result means that missing tendon cells in *dar1* KD disc are not cells that have lost their sr-gal4-positive fate. We therefore conclude that dar1 is not required to maintain *stripe* expression in developing tendon cells.

**Fig. 7.**
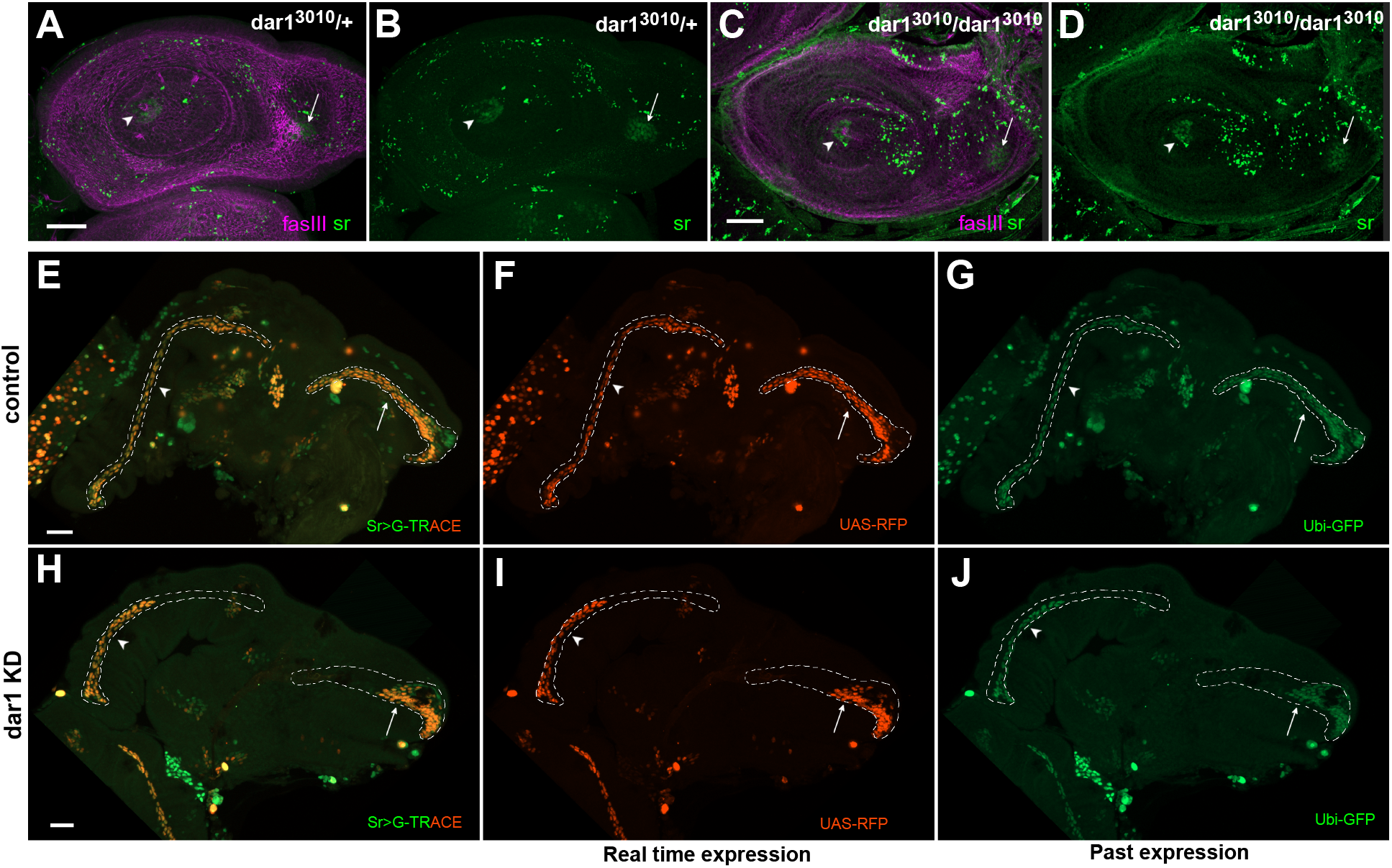
dar1 is not required to initiate or maintain *stripe* expression. (A-D) Confocal sections of L3 leg discs immunostained with anti-sr (green) and anti-fasIII (magenta). (A-B) In *dar1*^*3010/+*^ heterozygous leg discs, *stripe* expression reveals tendon cell precursors in tarsal segments (arrowhead) and in dorsal femur (arrow). (C-D) In the leg disc of rare *dar1*^*3010/3010*^ homozygous mutant escapers, *stripe* expression is still observed in tarsal segments (arrowhead) and dorsal femur (arrow). (E-J) G-TRACE in sr-gal4,dar1^3010/+^ (E-G) and sr-gal4,dar1^3010/+^>UAS-dar1RNAi (H-J) leg discs at 5h APF. RFP (red) staining shows current sr-gal4 expression (F, I) and GFP (green) reveals past expression (G, J). (E,H) merged G-TRACE expression. Dashed lines outline tarsal lt (arrowheads) and tilt in dorsal femur (arrows). At this stage, lt and tilt have deeply invaginated into the developing leg disc in the control (E-G), with most of the tendon cells having maintained sr-gal4 expression (red and green staining). By contrast, in *dar1* KD leg discs (H-I), the same tendons appear much shorter with both current sr-gal4 expression (red) and past sr-gal4 expression (green), indicating that shortening of those tendons is not due to premature loss of sr-gal4 expression in “missing” cells (prefigured by extension of dashed lines). Scale bar 30 µm.

Taken together, our results show that dar1 is required for a proper morphogenesis of the long internal leg tendon. The default of elongation observed in the *dar1* KD context is correlated with both actin polymerization defect and a dramatic decrease in sr-positive cell numbers, which cannot be explained by cell death, proliferation defect or loss of *stripe* expression. This leads us to propose a model in which the recruitment of new sr-positive cells is mediated by the tendon morphogenesis itself, and which supports a non-autonomous role of dar1 in tendon cell specification.

## Discussion and Perspectives

In *Drosophila* during metamorphosis, epithelium-derived progenitors of tendon legs undergo critical morphogenetic changes to adopt a unique tube-like structure for muscle attachment sites. Long tendon morphogenesis is characterized by several steps considered hallmarks of canonical tubulogenesis, such as cell apical constriction and cell invagination, followed by collective cell migration (Girdler and Röper, 2014; Hayashi and Kondo, 2018; Maruyama and Andrew, 2012). Another mark of tubulogenesis is that developing long tendons form an apical lumen and display F-actin-rich protrusions at the basal membrane of migrating cells (Laddada et al., 2019). Our newly-generated transcriptomic data, specific to leg tendon precursors, are consistent with these observations. These data highlight the expression of genes related to cell-shape rearrangement and cytoskeleton regulation, echoing the deep cell rearrangement of these epithelium-derived cells. Corroborating the importance of the tubulogenic process in leg tendon morphogenesis, GO analysis highlighted GO terms such as tube size regulation and morphogenesis. We also validated the involvement of the tubulogenic process by recapitulating the phenotypes observed in other systems, such as the tube length defects observed when *serpentine* and *vermiform* expressions are knocked down in the tracheal system (Luschnig et al., 2006).

To pinpoint critical transcriptional regulators of leg tendon development, we performed a short lethality and climbing RNAi screen. Our gene preselection, based on our transcriptomic data, was especially helpful: nearly 65% of these candidates induced locomotion defect and/or lethality when downregulated in tendon cells, even though RNAi knockdown efficiency is often limited by residual gene expression (Perkins et al., 2015). In comparison, a previous unbiased RNAi screen targeting 1384 genes to uncover genes involved in flight muscle attachment sites produced only about 1.5% positive candidates (Tiwari et al., 2015). Thus, the efficiency of our RNAi screen also emphasizes our RNA-seq data reliability.

Among the newly-identified candidates, we found that tendon expression of *dar1* and *CG9650* (data not shown) was restricted to the appendicular long tendon and was not found in other tendon precursors that do not form long internal structures (tendons of flight muscles or larval muscle tendon), making these genes excellent candidates for the specific development of long tendons. Correlating climbing defects, the LOF of these two genes impaired the formation of the leg long tendon. Whereas in *CG9650* KD tendons shaped correctly and later disrupted (data not shown), *dar1* KD led to a clear-cut tendon phenotype by the end of larval stages and early metamorphosis when long tendons are specified and elongate. Our further analysis of *dar1* function showed a critical role for this Krüppel-like factor in the elongation process, highlighting its requirement in actin cytoskeleton organization and specifically in the formation of actin-rich filopodia. Although there is no clear evidence for a direct role of filopodia in the tubulogenesis process, migrating cells form two types of actin-based protrusions, the lamellipodia, involved in cell motility, and the filopodia, which may sense and interact with the surrounding environment (Girdler and Röper, 2014; Okenve-Ramos and Llimargas, 2014; Svitkina et al., 2003). Interestingly, *dar1* has been shown to regulate the dendritic microtubule cytoskeleton by suppressing the expression of the microtubule-severing protein spastin (Ye et al., 2011), which we found depleted in our RNAseq data. Although we could not clearly elucidate whether manipulating spastin activity could indirectly affect filopodia formation in the developing tendon, reciprocal interactions between MT and actin networks is a well-established paradigm (Mohan and John, 2015; Rodriguez et al., 2003). For instance, the growth cone in neuronal cells depends on coordinated interactions between MTs and actin filaments (Bearce et al., 2015; Geraldo and Gordon-Weeks, 2009; Schaefer et al., 2002). We therefore cannot exclude the possibility that defects in actin organization observed upon dar1 downregulation are due to alterations of MT assembly and/or MT-actin network coordination. Supporting this last hypothesis, comparison of gene expression profiles between midgut cells expressing *dar1* RNAi and overexpressing *dar1* showed a deregulation of *DAAM* expression (Wu et al., 2018). *DAAM* encodes a formin known to link and coordinate actin and microtubule network dynamics (Szikora et al., 2017). Though to a lesser extent than *dar1* KD, our preliminary experiments designed to manipulate DAAM expression or activity also altered the formation of long tendon filopodia (not shown). It would thus be of interest to unravel a potential regulation of dar1 on DAAM, or more broadly, to perform a dar1 genetic screen interaction with actin and microtubule modulators.

Concomitantly to the default in long tendon growth and actin cytoskeleton organization, we demonstrated dar1 involvement in specification/recruitment of new sr-positive cells. First, we found that additional tendon progenitors were recruited, after initial expression of *stripe* in L3, contributing to the development of long tendons during their elongation in a normal context. Our finding that *dar1* KD led to a reduction of the number of sr-positive tendon progenitors and shortened tendons without affecting proliferation or increasing apoptosis suggested that *dar1* might control *stripe* expression (by inducing or maintaining its expression). However, our lineage and epistasis experiments indicated instead that dar1 acted downstream of stripe and that stripe was necessary for inducing *dar1* expression. We can thus infer that stripe regulates tendon morphogenesis through *dar1* activation, which is in turn responsible for the recruitment/specification of new sr-positive cells in a non-autonomous mechanism (our model **Fig. 8**). Organ morphogenesis often correlates with cell-fate specification, and numerous studies in the last decade have shown how multicellular morphogenetic events can feed back into gene regulatory pathways to specify cell fate (Chan et al., 2017). Thus during the 3D changes of a pseudo-stratified epithelium, the polarity and cytoskeleton rearrangements of the engaged cells are sensed by the neighboring cells through mechanotransduction. For instance, tissue deformations caused by germ-band extension upregulate the expression of the transcription factor twist via nuclear translocation of β-catenin (Desprat et al., 2008). We had previously shown that the Notch pathway is required for the initial expression of *stripe* in L3 stages (Laddada et al., 2019). However, conditional depletion of Notch activity at a later time point (early metamorphosis) did not appear to alter dorsal femur long tendon formation, whereas we show here that recruitment of additional sr-positive tendon precursors is still ongoing at this time. It is tempting to speculate that morphological changes, partly driven by *dar1*, could be mechanotransduced to the following cells and activate *stripe* expression in an alternative Notch-independent way. Alternatively, local reorganization of the leg disc epithelium by the primary invagination of first specified sr-positive cells could also play a role by exposing the surrounding cells to local signals. Kesavan et al. have demonstrated the role of CDC42-mediated tubulogenesis as a driver of non-cell-autonomous cell fate specification in pancreatic tubulogenesis (Kesavan et al., 2009). They showed how this small RhoGTPase, through its role in tube formation as an actin nucleator, provided the correct microenvironment for proper cell specification. Our own results also suggest that regulators of cytoskeletal dynamics and cellular rearrangement during tube formation can have an impact on the specification of tendon progenitors. Further work is required to determine to what extent dar1 transcriptional activity could regulate major actors of cytoskeleton remodeling and how spatial rearrangement of epithelial cells could provide a permissive microenvironment for a specifying signal that remains to be identified.

**Fig. 8.**
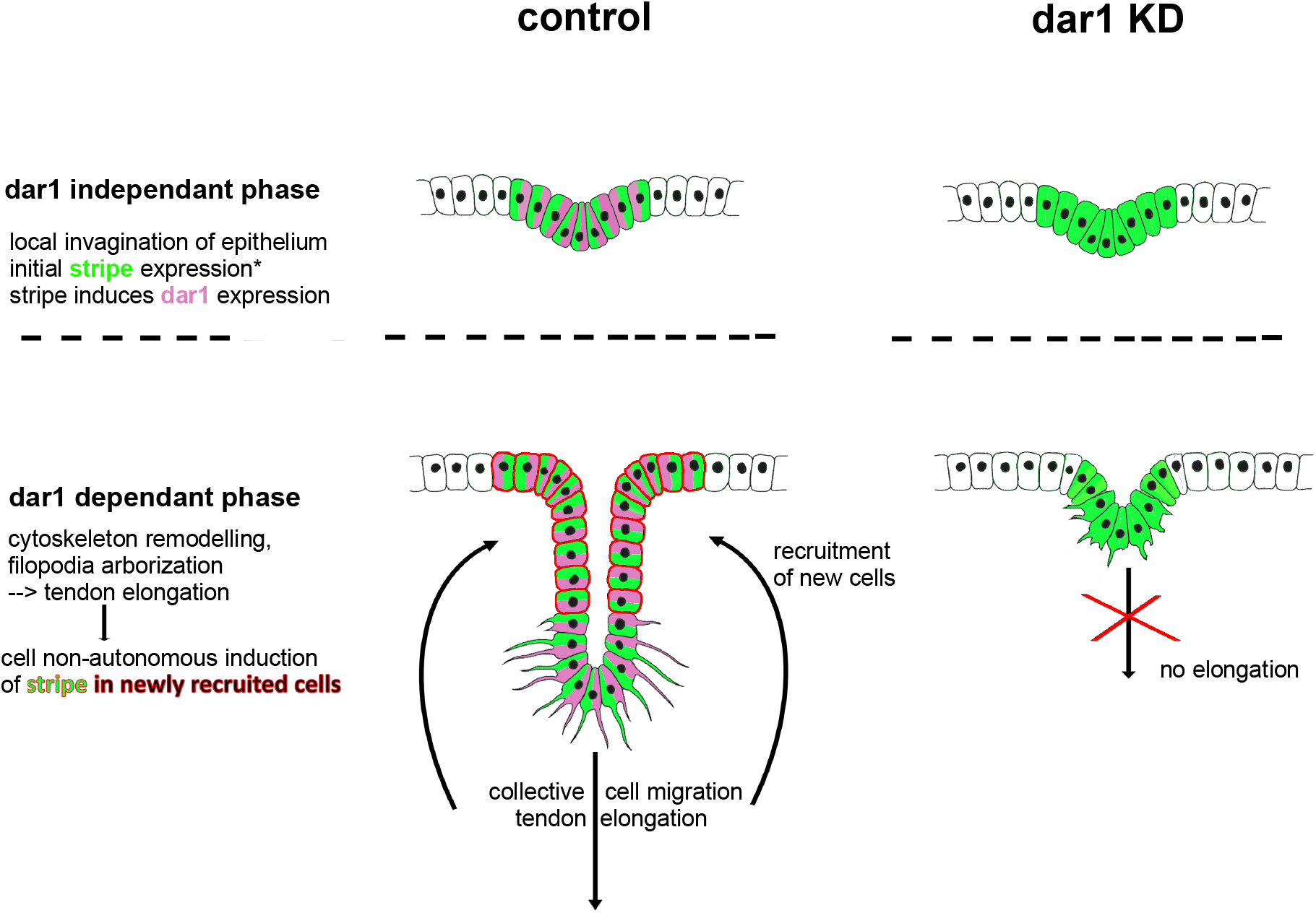
Model depicting dar1 function in the tubulogenic process of long tendon development. **dar1-independent phase:** in the first step of long tendon morphogenesis, Notch and odd are responsible for epithelium folding (invagination). During this phase, Notch is also responsible for first tendon progenitor commitment by inducing *stripe* expression in a few epithelial cells (Laddada et al. 2019): stripe is then required to induce *dar1* expression. **dar1-dependant phase:** during this second phase, dar1 regulates cytoskeleton remodelling and filopodia formation to promote collective cell migration and tendon elongation. In turn, the pulling mechanical forces could enable the recruitment of new sr-positive cells. In the absence of dar1, epithelium folding is not affected, but tendon elongation is aborted and there is no recruitment of new tendon (sr-positive) progenitors.

Although animal appendages have long been seen as non-homologous structures, there is growing molecular evidence that limbs of vertebrates and insects could arise from an ancestral appendage developmental program (Pueyo and Couso, 2005). Recently, Tarazona et al. reported that genes and signaling pathways that guide the development of both vertebrates and arthropods also control the development of cephalopod mollusk arms and tentacles (Tarazona et al., 2019). Their results thus strongly suggest that bilaterian appendages evolved by parallel activation of a genetic program that was present in a common ancestor. Whether the development of limb internal structures follows a similar conserved path is less well-documented. We have previously shown that the key regulator of vertebrate appendicular myogenesis, the *lb/Lbx1* gene, is also required for proper leg muscle identity in *Drosophila* (Maqbool et al., 2006). Strikingly, Huang et al. have shown that long tendons in the mouse form by a rapid elongation of the tendon in parallel with skeletal growth, and that this elongation is fueled by the recruitment of new mesenchymal progenitors (Huang et al., 2019). This recruitment is dependent on the transcription factor Scleraxis (Scx), which appears to be unnecessary for first tendon anchoring but is subsequently needed for the recruitment of new progenitors during tendon elongation. Although the *Scx* ortholog has not yet been identified in *Drosophila*, our study identified transcription factor dar1 as a specific factor of elongating tendons in *Drosophila*. Interestingly, its mammalian counterparts KLF5 and KLF4 have also been identified in two independent transcriptomic analysis of mouse limb tendon cells (Havis et al., 2014; Liu et al., 2015) and expression of KLFs have been reported in connective tissue surrounding tendon cells in the chicken (Orgeur et al., 2018). More strikingly, a preprint study (Kult et al., 2020) suggests that tendon-to-bone attachment cells have a bi-fated origin, with the ability to activate a combination of chondrogenic and tenogenic transcriptomes, and those authors identified KLF2/4 as central regulators of these unique bi-fated cells in vertebrates. Our work shows that *Drosophila* dar1/KLF-positive long tendons share a common origin with joint cells that connect exoskeletal elements of the leg. Thus, our and other most recent findings suggest an evolutionary conserved function of KLFs in the complex integration of the musculoskeletal system of the limb.

## Materials and Methods

The following *Drosophila* stocks were used: R10H12-gal4 (Pfeiffer et al., 2008 BDSC 48278), UAS-mCherryCAAX (BDSC 59021), UAS-mCherryNLS (BDSC 38425), enhancer trap lines sr-gal4^md710^ (Usui et al., 2004, BDSC 2663), UAS-Lifeact.GFP (BDSC 35544), UAS-dar1RNAi (BDCS 31987), dar1^3010^ (BDSC 65269), UAS-GC3Ai (Schott et al. 2017, BDSC 84346), UAS-serpRNAi (BDSC 63556), UAS-vermRNAi (BDSC 57188), UAS-lolalRNAi (BDSC 35722), UAS-Dicer2 (BDSC 24650), UAS-SrDN (gift from Vorbrüggen and Jäckle, 1997) and G-TRACE line (Evans et al., 2009, BDSC 28280).

### Cell sorting and RNA extraction

The fluorescence-activated cell sorting (FACS) protocol was adapted from (Harzer et al., 2013). Briefly, approximately 50 UAS-Lifeactin.GFP;sr-gal4/TM6b,tb white pupae (0h APF) were dissected on ice to collect 250–300 leg imaginal discs in M3 (S3652 Sigma Aldrich) complemented medium. Leg disc cells were dissociated in collagenase (P4762-Sigma Aldrich) and papain (C267-Sigma Aldrich) solution for 1h at 30°C, 300 rpm (Thermomixer Eppendorf), with additional mechanical stirring. After filtering, we used a FACSAriaTM device (4°C, 20 psi, nozzle Ø 100 µm) to first sort a batch of cells (roughly 50,000 cells per sample) independently of fluorescence, which we used as input (IP) and we then sorted the tendon cell (GFP+) population based on GFP fluorescence. Samples were directly collected in Trizol reagent (Invitrogen, 15596026). RNA extraction was performed with Zymo Quick-RNA microprep kit (R1051), sample RNA quality and quantity were estimated with QuBit (ThermoFisher, Qubit RNA HS Assay Kit, Q32852) and BioAnalyzer (Agilent, Agilent RNA 6000 Nano Kit 5067-1511), and sample specificity was analyzed by qPCR. Samples were stored at −80°C. For the detailed protocol, see Supplementary Material.

### RNAseq and transcriptomic data analysis

Extracted total RNA from GFP+ and IP cells from three different replicates were sent to Heidelberg genomic platform EMBL for high-throughput mRNA sequencing (NextSeq 500/Illumina). They generated mRNA libraries with NEB RNA Ultra kit (New England Biolabs E7770L) and dToligos probe were used to target mRNA for cDNA synthesis. Single-end multiplex was performed. Bioinformatic analysis was performed by Dr Yoan Renaud using FastQC and Bowtie2 software (reference genome Dm6). FPKM and differential expression between IP and GFP+ samples were determined using R script DEseq2 software. Sample correlations are given in Supplementary Material. Differential expression was performed using R-package DEseq2. Fold change (FC) between GFP+ and IP samples was computed using their normalized raw counts.

### *in vivo* RNAi screen

53 UAS-RNAi lines from two separate stock centres (VDRC and BDSC, see Supplemental Table 2) were crossed with UAS-dicer2, UAS-CAAXmcherry; sr-gal4/TM6b,tb,hu line and screened for lethality and/or climbing defect. 5–10 males carrying UAS-RNAi were mated with 15–20 females. Crosses and egg laying were performed at 25°C; 48h after egg laying, larvae were transferred at 28°C.

Embryonic or early larval stage lethality rate was measured by counting the number of nontubby over tubby third instar larvae. Percentage of lethality was calculated by applying the formula (1 - (number of nontubby larvae)/(number of tubby larvae)) × 100. Crosses were considered as positive for embryonic (early larval stage) lethality when lethality was higher than the 20% arbitrarily chosen threshold. To assess the percentage of lethality during metamorphosis, we applied the formula (1 - (the number of nontubby, nonhumeral emerging adult flies)/(total number of initial non tubby pupae)) × 100. Crosses with a higher score than 20% were considered positive candidates for metamorphosis lethality.

When sr-gal4>UAS-RNAi adult flies survived, we performed a climbing test adapted from the RING (Rapid Iterative Negative Geotaxis) assay published by Gargano et al. (Gargano et al., 2005). Up to ten young nonbalanced flies (48h old) were transferred to an empty vial and allowed to recover at room temperature for at least 2h. The vial was then rapped sharply on a table three times in rapid succession to initiate negative geotaxis responses. The flies’ positions in the vial were digitally captured after 5 s and we calculated the percentage of flies remaining in the bottom third of the vial. For each cross, two replicates (two vials of ten flies) were measured three times. We calculated the mean of these six trials. When this mean was equal to or greater than 30%, the corresponding cross was scored as “climbing defect”.

RNAis directed against *stripe* (100% embryonic lethality) and mcherry (0% embryonic lethality, 5% metamorphosis lethality, 15% climbing defect) were considered as positive and negative controls, respectively.

### Immuno-histochemistry, cryosections and confocal microscopy

The following primary antibodies were used: guinea pig anti-Dar1 (1:250, courtesy of B. Ye), guinea pig anti-stripe (1:500, from T. Volk), mouse anti-Fas III (1:500, Developmental Studies Hybridoma Bank [DSHB]), rabbit anti-Twist (1:500, our lab), mouse anti-Dlg (1:500, DSHB), chicken anti-GFP (1:500, Abcam), rat anti-DE-cadherin (1:500, DHSB), rabbit anti-Dcp1 (1:100, Cell Signaling), rabbit anti-pH3 (1:1000, Invitrogen), and anti-mcherry (rabbit, Abcam). Muscle fibers were visualized using cy3 or cy5-conjugated phalloidin (1:1000, Invitrogen) and nuclei using DAPI (ThermoFisher D1306). Secondary antibodies (Jackson) anti-rabbit, anti-guinea pig, anti-chicken and anti-mouse conjugated to Alexa488, cy3 or cy5 fluorochromes were used (1:500). Immunohistochemistry experiments were performed on samples fixed in 4% paraformaldehyde (PFA) for 20 min, rinsed in 0.1% PBS-Triton and blocked in 10% horse serum before immunostaining with primary antibodies (at 4°C, overnight). The samples were then rinsed and incubated with appropriate secondary antibody (at room temperature, 1h). For cryosections, we dissected adult leg and fixed them from 40 min to 1h in 4% PFA. Fixed samples were then incubated in sucrose solution (30%) at 4°C overnight. Adult legs were then laid on the bottom of a plastic well and carefully covered with Neg-50TM gel (Richard-Allan Scientific). The preparation was frozen at −80°C to set the gel. Sections were cut with a cryostat at 4°C (thickness 18–20 um). The sliced samples were stained with phalloidin before imaging. Immunostaining was visualized on an inverted SP8 Leica confocal microscope, and images were analyzed with Imaris 7.6.5 software.

### Tendon length measurement

Tendon size was determined by normalizing the length of the tendon over the whole size of the imaginal disc. For this purpose, we used the bounding-box function of the Imaris software to semi-automatically generate a 3D mesh of the imaginal disc (using FasIII staining). The disc was viewed as an ellipse with major axis (length) *X* and minor axis (width) *Y*. Discs were considered as flat objects, since at the developmental stages of analysis the thickness (*Z*) dimension has no significant impact on the measurement. From the ellipse-disc we computed the radius *R* of each corresponding disc (*R* =√ *X* ×*Y*/2). The lengths of tendons were measured using the function *MeasurementPoint* by manually following their pathways. Finally, we normalized the tendon length by dividing the length by the disc radius *R*. Statistical analyses were carried out using Prism software with an analysis of variance (ANOVA) between the different conditions and the different stages. Statistical significance between control and knockdown conditions was estimated using the Bonferroni test.

### Counting cell number in developing tendon

0h and 5h APF leg discs from sr-gal4,dar1^3010/+^>UAS-mcherryNLS and sr-gal4,dar1^3010/+^>UAS-mcherryNLS,UAS-dar1RNAi pupae were dissected and stained with DAPI for 1h. 0.5 µm confocal sections of long tendon of tarsi and dorsal tendon in femur (tilt) were performed. Images were processed with Imaris software and numbers of mcherryNLS and DAPI-positive cells per tendon were determined automatically using the function *Spots*. The automatic cell counting was subject to manual adjustement.

Statistical significance was estimated using the Bonferroni or Welch test.

